# Polyphyletic origins and lineage-specific expansion of chemosensory systems in phylum Actinomycetota

**DOI:** 10.64898/2026.11.03.764138

**Authors:** Sri Nithya S, Sherin Brighty, Gaurav Sharma

## Abstract

Phylum Actinomycetota (syn. Actinobacteriota or Actinobacteria), one of the largest well-studied bacterial phyla, is renowned for complex morphology, non-motility, ecological versatility, and secondary metabolite production. Chemosensory systems (CSS) are among the most sophisticated prokaryotic signal transduction architectures, yet their evolutionary distribution in this phylum remains poorly understood. Here, a comparative genomic analysis of 693 high-quality genomes spanning the phylum resolves the diversity, distribution, and evolutionary trajectories of CSS. Despite extensive genomic complexity and ecological versatility, only 6.9% of genomes encode complete CSS clusters, confined to a subset of lineages within a few classes, indicating lineage-specific retention or loss. In contrast, two-component systems (TCS) are ubiquitously abundant, with hybrid kinases enriched in CSS-deficient classes, suggesting functional compensation for the absence of dedicated chemosensory pathways. Basal lineages retain simplified CSS (F1 or F5), whereas Actinomycetes show lineage-specific expansion into multiple classes (F5, F8, F9, and ACF), often co-occurring, indicative of enhanced sensory complexity. The phylogeny of CheA homologs shows a polyphyletic distribution, supporting a mosaic evolutionary history shaped by multiple independent acquisitions, diversification, and gene loss. Collectively, our findings show that, unlike motile bacteria, Actinomycetota members rely predominantly on TCS-driven regulatory networks, with CSS retained only in select lineages where ecological demands require enhanced environmental sensing. This work provides a phylum-wide evolutionary framework for the emergence, diversification, and functional specialization of CSS in non-motile bacteria.

**Importance:** Bacteria rely on sensory signal transduction systems to detect and respond to environmental changes. Chemosensory systems typically control movement along with alternate cellular functions, but their role in non-motile phylum Actinomycetota has remained unclear. By analyzing approximately seven hundred Actinobacterial genomes, we report that these systems are rare and restricted to specific lineages, suggesting they are retained only where they confer ecological advantages. In contrast, most Actinomycetota depend on alternative signaling pathways, particularly two-component systems, for environmental adaptation. These findings highlight how different bacterial groups evolve distinct strategies for sensing and responding to their surroundings, providing new insights into microbial evolution, ecological specialization, and the emergence of complex adaptive traits.

## Introduction

Bacterial signal transduction systems enable these tiny prokaryotes to perceive and integrate environmental stimuli and coordinate adaptive responses governing chemotaxis, virulence, biofilm formation, antibiotic resistance, metabolic regulation, and interspecies communication (Krell et al., 2010). Based on their modular architecture, these systems are broadly classified into one-component systems (OCS), two-component systems (TCS), and chemosensory systems (CSS) (Ulrich et al., 2005; Galperin, 2010). OCS represents the most ancient and abundant class, comprising a single polypeptide that integrates both sensory input and regulatory output domains, with many characterized transcription factors belonging to this category (Ulrich et al., 2005). TCS, the predominant bacterial signaling mechanism, employs histidine kinase-mediated phosphorylation of a response regulator to modulate cellular outputs (Stock et al., 2000; Gao & Stock, 2009).

CSS represents a multi-protein microbial (bacterial and archaeal) sensory architecture, in which all constituent proteins are functionally specialized and operate as integrated signaling assemblies (Bi & Sourjik, 2018). The core chemosensory pathway is composed of methyl-accepting chemotaxis proteins (MCPs), which function as transmembrane chemoreceptors; CheA, a histidine kinase that is activated upon MCP conformational changes mediated by ligand binding; and CheY, a response regulator whose phosphorylated form (CheY-P) interacts with the flagellar motor switch complex to modulate rotational direction (Sourjik & Berg, 2002; Hazelbauer et al., 2008). Signal gain and receptor sensitivity are regulated by the methyltransferase CheR and the methylesterase CheB, which together implement a sensory adaptation mechanism through reversible methylation and demethylation of conserved glutamate residues in MCP cytoplasmic domains, respectively (Stock J et al., 1978; Kentner & Sourjik, 2006). CheW, a scaffold protein bearing a PH-like domain, mediates the formation of ternary receptor–kinase complexes at the cell pole, facilitating cooperative signaling and signal amplification (Briegel et al., 2012; Cassidy et al., 2020). CheZ accelerates CheY-P dephosphorylation to reset the signaling circuit and enable rapid adaptation (Zhao et al., 2002). Together, these components form a highly cooperative, ultrasensitive signaling network capable of detecting attractant/repellants concentration changes as small as 3nM over a background spanning five orders of magnitude (Sourjik & Berg, 2002).

Three different types of CSS classes are known, flagella-mediated motility systems (Fla) for flagellar motility, type IV pili-based motility systems (Tfp) for pilus mediated motility, and alternative cellular function systems (ACF), which regulates non-motility processes including cell cycle progression, biofilm formation, exopolysaccharide biosynthesis, and, paradoxically, even flagellar biosynthesis itself (He & Bauer, 2014; Munar-Palmer et al., 2024). The Fla class has undergone remarkable diversification into 17 phylogenetically distinct subclasses (F1–F17), whereas Tfp and ACF each constitute a single subclass, reflecting the evolutionary specialization and functional constraint of flagellar-based chemosensory architectures (Colin et al., 2021). Evolutionary analyses suggest that ACF systems may represent an ancestral state among CSS, given their comparatively simple protein composition and functional parallels to MAC (Methyl-accepting Chemotaxis-like) regulatory systems (Wuichet & Zhulin, 2010). As motility functions evolved, these ancestral CSSs likely diversified and disseminated across diverse bacterial and archaeal lineages to regulate diverse physiological functions via ACFs (Zhulin, 2001; Wuichet & Zhulin, 2010).

Chemosensory systems have been widely characterized in several model organisms, such as *Escherichia coli* and *Bacillus subtilis* each possess a well-characterized chemotaxis system, whereas *Pseudomonas aeruginosa* PAO1 encodes four CSSs (F6, F7, ACF, and Tfp) and *Azospirillum sp.* B510 contains six CSSs, comprising F5, F7, F8, F9, and two ACF systems. The more complex sensory repertoires of *Myxococcus xanthus* DK1622 and *Cystobacter fuscus* DSM2262 include eight and 12 CSSs, respectively; the latter comprises F1, F10, three F8 systems, and seven ACF systems (Sharma G et al., 2018). Such variation in CSS abundance, class composition, and regulatory architecture likely reflects lineage-specific adaptations to diverse ecological and physiological demands, suggesting the evolutionary significance of chemosensory signaling in microbial environmental adaptation (Alexander & Zhulin, 2007)

This variation raises an important evolutionary question concerning bacteria that possess complex genomes and elaborate developmental or ecological lifestyles but lack canonical motility-associated sensory systems. The phylum Actinomycetota, formerly referred to as Actinobacteria or Actinobacteriota, provides an appropriate context in which to investigate this question. Actinomycetota is a phylogenetically diverse and ecologically widespread bacterial phylum comprising predominantly Gram-positive organisms with characteristically high genomic GC content and considerable variation in genome size (Seshadri et al., 2022). Its members occupy diverse environments, including terrestrial soils, rhizospheres, aquatic habitats, and host-associated niches, and exhibit substantial metabolic versatility (Hashim et al., 2026). Several lineages possess large genomes, ranging from ∼0.5 Mbp (in obligate intracellular pathogens such as *Tropheryma whipplei*) to >15 Mbp (in morphologically complex genera such as Streptomyces) (Barka et al., 2016; Bentley et al., 2002), enriched in regulatory genes and biosynthetic gene clusters, contributing to their capacity for secondary metabolite production, environmental adaptation, and morphological differentiation (Barka et al., 2016; Jorgensen et al., 2024; Hashim et al., 2026). The genus *Streptomyces*, in particular, is recognized for its complex developmental biology and extensive biosynthetic potential, including the production of antibiotics, antifungals, and other bioactive compounds (Antoraz et al., 2020; Barka et al., 2016). Furthermore, *Streptomyces* species utilize volatile organic compounds as airborne signals to regulate secondary metabolism and activate silent pathways in neighboring Actinobacteria (Cuervo et al., 2024). Actinomycetota therefore combines substantial genomic and ecological complexity with predominantly non-motile lifestyles, making it an informative system for examining the evolutionary distribution of sensory signaling architectures.

However, despite these hallmarks of genomic and phenotypic complexity, Actinomycetota does not necessarily rely on the same sensory strategies as other bacteria with complex lifestyles. In Actinomycetota, chemosensory systems (CSS) are noticeably sparse or altogether absent in most members, with environmental sensing instead dominated by a markedly expanded repertoire of two-component systems (TCS) (Wadach et al., 2025). The most known model organism, *Streptomyces coelicolor* A3(2) encodes 109 histidine kinase, 86 response regulators and only two hybrid histidine kinases alongside an extraordinary diversity of 58 extracytoplasmic function (ECF) sigma factors, enabling transcriptional integration of environmental signals without reliance on motility-coupled chemosensory circuits (Sanchez de la Nieta R et al., 2022; Hutchings et al., 2004; Paget et al., 2002). Most existing signal transduction studies in actinobacteria have focused on OCS and TCS-mediated regulation of antibiotic biosynthesis and morphological differentiation in *Streptomyces* (MCLean TC et al., 2026; Hutchings et al., 2004; Paget et al., 2002). A few studies have also focused on undertaking a phylum-wide, comparative assessment of two signal transduction classes, OCS and TCS (Sanchez de la Nieta et al., 2022; Cruz-Bautista et al., 2023; Gongerowska-Jac et al., 2021). A useful comparative perspective is provided by the phylum Myxococcota, whose members exhibit similar large genome size and complex multicellular behaviors via predation, social gliding, fruiting-body formation, and developmental transitions, through extensive regulatory systems that include both TCS and CSS (Muñoz-Dorado et al., 2016; Shi et al., 2008; Barry et al., 2006). *Myxococcus xanthus* DK1622, for example, possesses a large repertoire of TCS components and multiple CSS classes associated with flagellar motility, type IV pili-mediated gliding, and alternative cellular processes (Sharma et al., 2018). The contrasting sensory architectures of Myxococcota and Actinomycetota provide an opportunity to examine how bacterial lineages with substantial genomic and ecological complexity have evolved different strategies for environmental sensing. Whereas Myxobacteria combine extensive CSS with other regulatory pathways, Actinomycetota may rely more heavily on TCS and other transcriptional regulatory mechanisms, with CSS retained only in particular evolutionary lineages. The extent and evolutionary basis of this contrast, however, have not been systematically resolved across Actinomycetota.

Overall, a comprehensive assessment of CSS across Actinomycetota is therefore necessary to determine how these systems are distributed across their major lineages and how their architectures have diversified during phylum evolution. Such an analysis can establish whether CSS are broadly conserved or phylogenetically restricted, identify lineages exhibiting expansion or reduction of specific CSS classes, and clarify the relationship between CSS diversity and the broader repertoire of signal transduction systems. Furthermore, examining the evolutionary relationships of conserved CSS components, such as CheA, may provide insights into the relative contributions of vertical inheritance, lineage-specific diversification, horizontal gene transfer, and gene loss to the present-day distribution of these systems.

Therefore, this study addresses this gap by presenting a comprehensive, genome-scale analysis of TCS and CSS distribution across a phylogenetically diverse set of 693 Actinomycetota genomes, taxonomically categorized into six classes, 29 orders, 64 families, 187 genera and 693 strains. By systematically cataloguing the repertoire and composition of signal transduction proteins across ecologically distinct members of Actinomycetota, spanning soil-dwelling, host-associated, aquatic, and morphologically complex lineages, we aim to resolve whether the CSS-stunted architecture observed in well-studied actinobacterial model organisms reflects a phylum-wide evolutionary constraint or a lineage-specific reduction associated with the loss of motility. Collectively, the findings presented here provide a foundational framework for understanding how regulatory CSS diversity and complexity is achieved and maintained in the absence of canonical chemosensory pathways, with broader implications for the evolution of non-motility-associated signal transduction across the bacterial domain.

## Materials and methods

### Data sources for comparative genome analysis

Actinomycetota genomes were selected from the prokaryotes section of the NCBI Genome FTP repository, restricted to reference and complete genomes deposited between the years 2007 and 2024. 88,957 Genome assemblies and annotations were downloaded using the NCBI Datasets command-line tool, which has 897 complete reference genomes. A total of 693 genomes were curated for analysis, with taxonomy mapped using the prokaryotes.txt file from the NCBI Taxonomy FTP resource. A comprehensive genome statistics table summarizes the taxonomy, assembly attributes, genome size, GC content, and CDS features for all genomes under study (Table S1). Ecological metadata was extracted using the FetchM Python module for a subset of organisms (Anik, 2025).

### Identification and categorization of the TCS/CSS proteins and clusters

The translated CDS files of all selected Actinomycetota genomes were mapped against the Pfam-A v34.0 database (Mistry J et al., 2021) using the HMMER suite (Finn et al., 2011) with an E-value cutoff of 1e^−5^ to identify conserved protein domains. The resulting hits were then analyzed using the hmmscan-parser.sh script to identify non-overlapping functional domains. To identify proteins involved in signal transduction pathways, particularly two-component systems (TCS) and CSS, Pfam domain annotations were cross-referenced with information from MiSTdb 4.0 (Gumerov et al., 2024). The typical domains corresponding to TCS proteins included Response_reg (PF00072), HATPase_c (PF02518), HATPase_c_2 (PF13581), HisKA (PF00512), HisKA_2 (PF07568), HisKA_3 (PF07730), and Hpt (PF01627). Similarly, characteristic domains associated with CSS proteins were identified as CheB_methylest (PF01339), CheC (PF04509), CheD (PF03975), CheR (PF01739), CheR_N (PF18676), CheW (PF01584), CheX (PF13690), CheZ (PF04344), and MCPsignal (PF00015). Domain profiles were used to classify and quantify signal transduction proteins across the genomes analyzed. CSS components were identified using an in-house pipeline following the MISTdb (Gumerov et al., 2024) for all 693 Actinomycetota genomes. An overview of TCS (Table S2) and CSS (Table S3) repertoire variation within phylum has been shown, which consists of species-level taxonomic information along with encoded proteins for diverse signal transduction categories and CSS classes.

### Phylogenetic analysis of housekeeping genes

The proteomes of all 693 Actinomycetota genomes were subjected to orthologous gene clustering using ProteinOrtho v6.3.4 (Lechner et al., 2011) to delineate single-copy orthologous genes conserved across the entire dataset. Only three universally conserved single copy orthologs were recovered from all genomes, corresponding to the core housekeeping genes (encoding the 50S ribosomal protein L16, elongation factor P, and 30S ribosomal protein S6), selected as phylogenetic markers based on their ubiquitous distribution and functional conservation across the phylum. The corresponding protein sequences were individually aligned using MUSCLE v5.1 (Edgar, 2022) employing the super5 algorithm, which is optimized for high-accuracy multiple sequence alignment of large datasets. The resulting individual alignments were subsequently concatenated into a single alignment block, which served as the input for phylogenetic inference using FastTree (Price et al., 2010), an approach that constructs approximately maximum-likelihood phylogenies with high computational efficiency. The resulting core housekeeping gene phylogeny was interactively visualized and taxonomically annotated using the Interactive Tree of Life (iTOL) platform (Letunic & Bork, 2024) to facilitate comparative phylogenomic interpretation.

### Modular representation of CSS

A phylogenetic tree was created based on the CheA sequence of seventy-one representative sequences of Actinomycetota species that contain CSS. The modular arrangement of CSS was evaluated in each resulting cluster of the phylogenetic tree to determine any structural similarities and differences related to types of flagellar chemosensory classes.

### Phylogenetic analysis of CSS

71 CheA homologs identified from 48 Actinomycetota species were analyzed using BLASTp v2.13. (Camacho et al., 2009) against the NCBI NR database (Sayers et al., 2025) with an E-value cutoff of 1e^-5^. Homologs identified based on >80% identity were extracted along with 900 non-redundant sequences from other bacterial phyla as outgroups. All sequences were aligned using MUSCLE v5.1 (super5 algorithm), and a phylogenetic tree was constructed using FastTree (Price et al., 2010). The phylogeny was visualized in iTOL (Letunic & Bork, 2024) to examine the evolutionary distribution and directions of Actinomycetota CheA proteins.

## Results

### Taxonomic, ecological, and genomic composition of the Actinomycetota dataset is dominated by class Actinomycetes

To characterize signal transduction diversity across the phylum Actinomycetota, we assembled a dataset of 693 complete reference genomes from the NCBI Genome database, selected from 897 complete reference genomes after filtering based on complete genome assemblies meeting stringent quality thresholds (Fig. 1A). Taxonomic assignments, assembly statistics, genome size, GC content, and CDS counts for each genome are provided in Supplementary Table S1. The dataset spans six classes: Acidimicrobiia, Actinomycetes, Coriobacteriia, Nitriliruptoria, Rubrobacteria, and Thermoleophilia (Fig. 1B). Actinomycetes accounted for 655 of the 693 genomes (94.5%), whereas the remaining five classes together contributed thirty-eight genomes (5.5%; Coriobacteriia, 19; Thermoleophilia, 7; Acidimicrobiia, 5; Rubrobacteria, 4; Nitriliruptoria, 3). This imbalance most likely reflects the over-representation of clinically and industrially important genera, such as Streptomyces, Mycobacterium, and Micromonospora, in public sequence databases (Barka et al., 2016), and it constrains confidence with which phylum-wide inferences can be drawn for the under-sampled classes. Within Actinomycetes, genomes were distributed across 29 orders (Fig. 1B), of which Mycobacteriales (n = 201), Micrococcales (n = 163), Kitasatosporales (n = 87), and Propionibacteriales (n = 53) were the most strongly represented.

**Figure 1:**
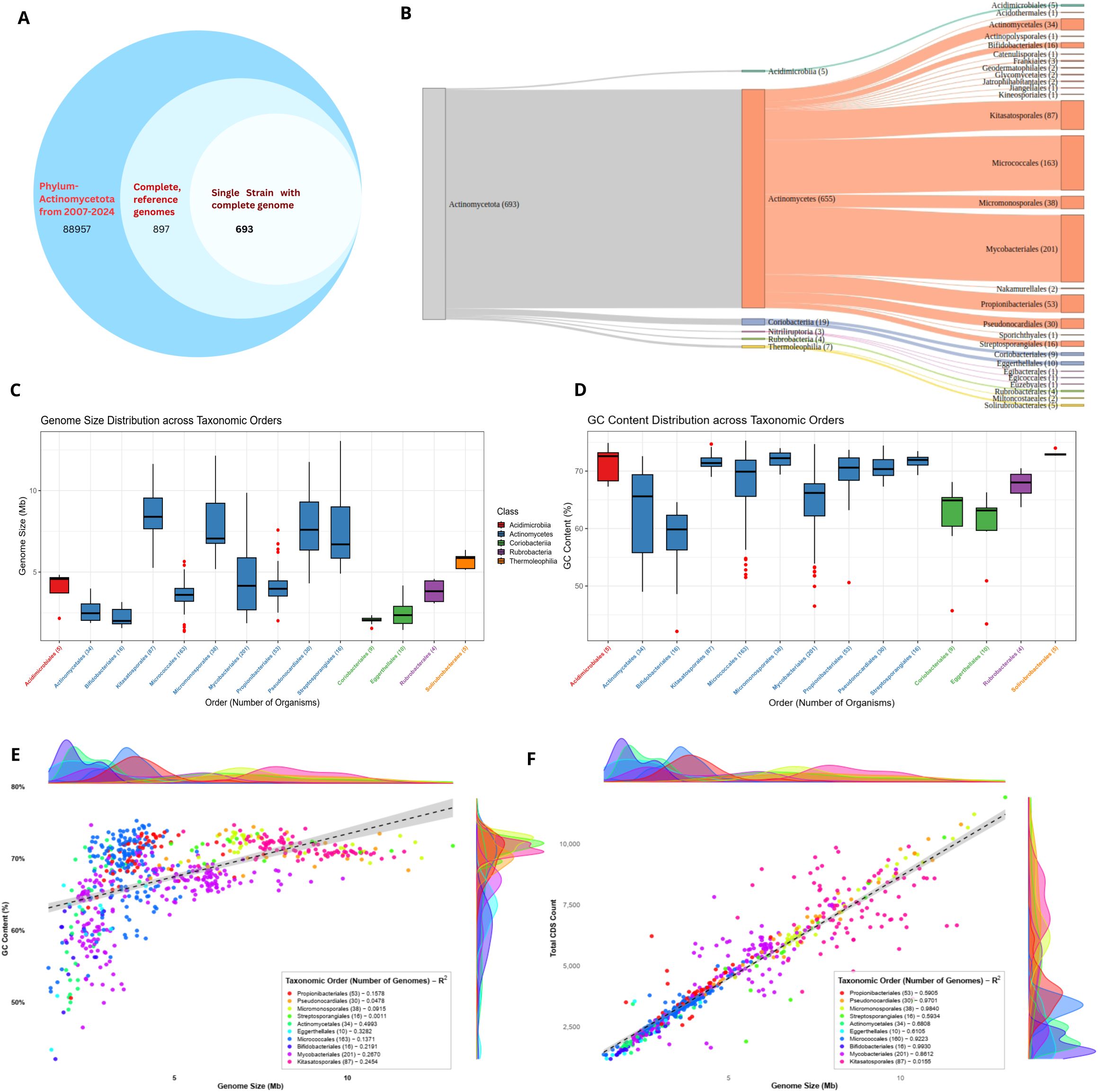
Dataset composition and genomic features across Actinomycetota: **(A)** Circular schematic illustrating the genome selection strategy, showing the progressive filtering of phylum Actinomycetota genomes from NCBI to obtain 693 high-quality representative complete genomes. (**B)** Sankey graph illustrates the hierarchical taxonomic flow from phylum to class and order, highlighting the asymmetric representation driven by Actinomycetes. (**C & D)** Genome size and GC content variation across taxonomic orders visualized as box plot, labelled with their respective taxonomical class. **(E)** Density scatter plot illustrates the weak association between genome size and GC content across taxonomic orders with at least three representative genomes, with the marginal density plots depicting the order-specific distribution patterns. **(F)** Density scatter plot depicts the strong positive relationship between genome size and the number of CDS across taxonomic orders with at least three representative genomes, with marginal density plots highlighting the order-specific distribution patterns.

Habitat information was available for 388 of the 693 genomes and was grouped into four categories: host-associated, terrestrial land, terrestrial water, and marine (Fig. S1; Table S1). Among these, Mycobacteriales, Micrococcales, Kitasatosporales, Propionibacteriales, and Pseudonocardiales included members from three or four habitat categories, suggesting generalist ecological strategies and broad environmental adaptability. By contrast, Bifidobacteriales and Eggerthellales were almost exclusively host-associated, which is consistent with niche specialization and the genome reduction typical of host-adapted bacteria (Toft & Andersson, 2010). Because roughly 44% of genomes lacked habitat annotation, these ecological patterns should be regarded as indicative rather than definitive.

Genome size and GC content varied considerably among classes and orders (Fig. 1C, D). Actinomycetes showed the widest genome size range (1.36-13.05 Mb; median, 4.21 Mb; mean, 5.02 Mb) and a median GC content of 69.2%. Within this class, genome size ranged from the compact genomes of orders such as Bifidobacteriales and Actinomycetales (median ∼2-2.5 Mb) to the large genomes of Kitasatosporales, Micromonosporales, and Streptosporangiales (medians of approximately 7 to 8.5 Mb), the former including host-associated taxa and the latter predominantly soil-dwelling genera, such as Streptomyces and Micromonospora (Jorgensen et al., 2024) (Fig. S2, Fig. S3). Coriobacteriia had the smallest genomes among the classes (median, 2.1 Mb), whereas Thermoleophilia combined moderately large genomes (median, 5.2 Mb) with the highest median GC content (73.0%), overall suggesting that GC enrichment in these lineages is maintained independently of genome size, potentially reflecting distinct mutational biases or DNA repair mechanisms rather than genome-size-linked selection (Hildebrand et al., 2010; Bohlin et al., 2010).

Correlation studies between genome size, GC content, and CDS count (Fig. 1E, F) revealed that genome size and GC content were moderately correlated across the dataset (Pearson’s r = 0.51). However, the strength of these associations differed among orders, and coefficients of determination within individual orders were generally low (R² ranging from near zero to approximately 0.5), with the highest values in Actinomycetales, Eggerthellales, and Mycobacteriales (Fig. S4, Fig. S5). These results indicate that GC content varies largely independently of genome size within most lineages, which suggests that lineage-specific factors, such as mutational bias, codon usage, and repair-pathway differences, rather than genome-size-linked selection shape nucleotide composition (Hildebrand et al., 2010; Bohlin et al., 2010; Foerstner et al., 2005). In contrast, Genome size scaled strongly and positively with total CDS count across the dataset (Pearson’s r = 0.94; Fig. 1F). Within orders, the fit was heterogeneous. Five of the ten orders shown had R² > 0.90, three orders were intermediate (Actinomycetales, Mycobacteriales, Eggerthellales), whereas two orders (Micrococcales 0.02, Kitasatosporales 0.02) did not show any good correlation, despite their large sample sizes (n = 163 and 87). Coding gene content is therefore the principal correlate of genome size in most lineages, which indicates that genome expansion is driven mainly by gain of protein-coding sequence rather than non-coding DNA. In Micrococcales and Kitasatosporales, however, CDS count is largely decoupled from genome size, which is consistent with lineage-specific differences in coding density or in non-coding content. The coding-driven pattern is in line with reports linking larger actinomycete genomes to expansion of biosynthetic, regulatory and signal-transduction gene families through vertical inheritance and horizontal gene transfer (Cimermancic et al., 2014; Huang et al., 2015).

Collectively, these results establish substantial intra-phylum variation in taxonomic representation, ecology, and genome architecture, with class Actinomycetes contributing most of this diversity and genome expansion proceeding primarily through gains in coding capacity. This genomic and ecological framework, spanning generalist, broadly distributed lineages to reduced, host-associated genomes, provides the taxonomic backdrop against which the distribution of signal transduction systems, examined in the following sections, can be interpreted.

### Two-component system proteins dominate signal transduction in Actinomycetota, including CSS-deficient classes

To establish a phylum-wide quantitative baseline of signal transduction capacity, all 693 Actinomycetota genomes were systematically screened for proteins belonging to TCS and CSS. Similar to previous studies (Ulrich et al., 2005), TCS-associated proteins represented 25.2% of all signal transduction proteins identified, encompassing orphan HKs and orphan RRs (both lacking the other cognate partner in the same protein), and canonical hybrid histidine kinase-response regulator (HK-RR) pairs where histidine kinase and receiver domains are fused within a single polypeptide (hybrid HKs, HHKs; hybrid RRs, HRRs). The prevalence of orphan regulators is noteworthy, as unpaired HKs and RRs have been proposed to participate in cross-regulatory phosphotransfer networks or to function as dedicated phosphate sinks that modulate the activity of cognate TCS pairs, a regulatory strategy documented in organisms with complex developmental programs (Laub & Goulian, 2007; Groban et al., 2009).

TCS component abundance was evaluated as both absolute copy numbers and genome-size-normalized counts (per Mbp) to account for the substantial genome size variation documented across the phylum (Fig. 2A-F). The phylum-wide median was 27 HKs and 26 RRs per genome, which indicates stoichiometric coupling of sensor kinases and response regulators consistent with the coordinated architecture and signaling balance described previously (Capra & Laub, 2012; Krell et al., 2010; Laub & Goulian, 2007). It must be noted that as Actinomycetes make up 94.5% of the dataset, these values largely reflect that class. Furthermore, the median total TCS count per genome was 52, scaling closely with genome size (r = 0.91). Median TCS density was 11-13 per Mbp in Actinomycetes, Coriobacteriia and Acidimicrobiia, and higher in Nitriliruptoria (16.8), Rubrobacteria (19.3) and Thermoleophilia (21.4). The compact genomes of the CSS-deficient classes therefore carry TCS repertoires that are proportionally as large as, or larger than, those of Actinomycetes.

**Figure 2:**
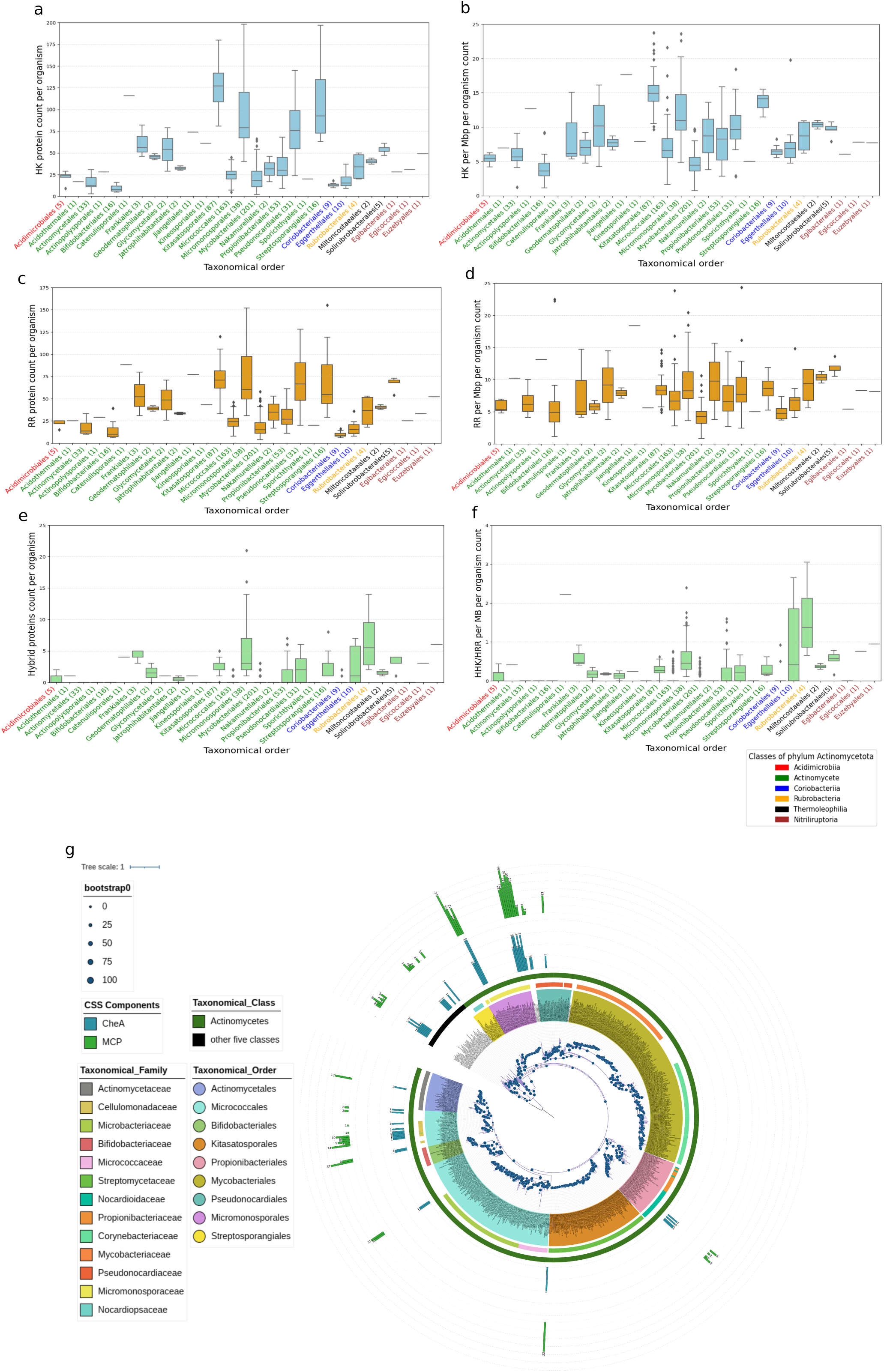
Distribution and phylogenetic mapping of signal transduction components across Actinomycetota. **(A, C, E)** Box plots represent the absolute distribution of HK, RR, and hybrid proteins, respectively, across six taxonomical classes at the order level. X-axis labels are color-coded by taxonomical class, with the number of genomes represented by each order indicated in parentheses. **(B, D, F)** Box plots depict the normalized distributions of HKs, RRs, and hybrid proteins per megabase of genome size, respectively, highlighting lineage-specific scaling of signaling components independent of genome size. (**G)** Rooted circular phylogeny of 693 Actinomycetota genomes reconstructed using three conserved core proteins, with *Bacillus subtilis* (Firmicutes) included as the outgroup. The three genes were aligned separately and concatenated to construct the core-gene phylogeny. The innermost colored ring denotes taxonomic order, followed by the outer rings representing family and class, respectively. CSS components are mapped onto the terminal branches, where CheA and MCP counts are displayed as radial bars in their corresponding colors.

Hybrid kinases (HHKs) and hybrid response regulators (HRRs), which are characteristic components of multi-step phosphorelays capable of integrating multiple input signals before committing to a regulatory output (Mo et al., 2022), were disproportionately enriched in the orders of non-chemosensory classes, most notably Eggerthellales within Coriobacteriia and Rubrobacterales within Rubrobacteria. The prominence of hybrid systems in these CSS-lacking classes is consistent with the hypothesis that phosphorelays may functionally substitute for the signal integration and temporal filtering properties normally provided by CSS in motile bacteria, enabling nuanced regulatory responses to environmental fluctuations without reliance on chemosensory machinery (Capra & Laub, 2012; Mo et al., 2022).

### Chemosensory systems are sparsely and non-randomly distributed across Actinomycetota, with the majority in Actinomycetes

Because a few individual Che proteins, e.g., stand-alone CheY homologs, or MCPs coupled to non-CSS outputs, can function outside canonical chemosensory pathways (Wuichet & Zhulin, 2010), we applied a conservative criterion: a genome was scored as encoding a CSS only if it carried at least one CheA homolog, the non-redundant histidine kinase of the pathway (Bi & Sourjik, 2018; Munar-Palmer et al., 2024). Applying this threshold, only forty-eight of the 693 genomes analyzed (6.9%) were identified as encoding a complete chemosensory system, cumulating to total seventy-one systems within 48 organisms (Table S2). CSS-encoding genomes were exclusively distributed across four of the six classes represented in the dataset, Acidimicrobiia, Actinomycetes, Nitriliruptoria, and Thermoleophilia, with Coriobacteriia and Rubrobacteria being entirely devoid of CSS, suggesting either ancestral CSS absence or complete secondary loss in these lineages (Wuichet & Zhulin, 2010).

To place this distribution in an evolutionary framework, we reconstructed a maximum-likelihood phylogeny of all 693 genomes from three concatenated single-copy orthologs (L16, EF-P, S6), rooted with *Bacillus subtilis* (Fig. 2G). The resulting topology recapitulated the major deep-branching divisions within Actinomycetota, with well-supported clades corresponding to established taxonomic orders, on which CheA and MCP distribution was mapped. CheA and MCP proteins were phylogenetically restricted to specific well-supported clades within Actinomycetes, with sporadic presence observed in several orders including, Actinomycetales (1/34), Micrococcales (14/163), Propionibacteriales (4/53), Micromonosporales (11/38), and complete absence across Mycobacteriales (201 genomes), Kitasatosporales (87 genomes), Streptosporangiales (16 organisms), and Bifidobacteriales (16 organisms) (Fig. 2G). The congruence between CheA distribution and phylogenetic topology, with CSS-encoding lineages clustering within specific clades rather than being randomly distributed across the tree, suggests that chemosensory capacity in Actinomycetota is the product of lineage-specific retention and elaboration following an ancestral CSS acquisition event, rather than widespread independent gain across the phylum. This phylogenetic signal is further consistent with the ecological partitioning observed, wherein soil-dwelling and aquatic orders with broader ecological ranges were disproportionately represented among CSS-encoding lineages, while obligate host-associated orders such as Bifidobacteriales (Fig S1), which undergo reductive genome evolution and face reduced selective pressure for environmental motility-sensing, exhibited complete CSS absence (Moran, 2002; Giovannoni et al., 2014). Together, these observations establish that the signal transduction architecture of Actinomycetota is defined by TCS dominance, with CSS representing a phylogenetically and ecologically constrained accessory feature confined to a minority of lineages that likely retain or have elaborate chemosensory capabilities in association with specific ecological demands.

### Lineage-specific expansion of chemosensory systems in class actinomycetes

Mapping CSS classes onto the core-gene phylogeny (Fig. 3A) and tabulating them by class (Fig. 3B-C) revealed a strong asymmetric distribution. Several early branching classes like *Coriobacteriia* and Rubrobacteria completely lacked CSS systems. Thermoleophilia, another basal group, exclusively encoded F1 systems, suggesting retention of a simple, conserved architecture. In contrast, *Acidimicrobiia* possessed only F5 systems, indicating an independent early divergence of this architecture outside Actinomycetes. Remarkably, the late-diverging class *Actinomycetes* contained representatives of F5 (37/42), F8 (2/2), F9 (8/8), and ACF systems (12/12) across a small number (36 genomes) out of 655 genomes. This dominant accumulation of F5, together with the emergence of the rare F8 and F9 systems exclusively within this lineage, confirms a major expansion of chemosensory capabilities during Actinomycetes evolution.

**Figure 3:**
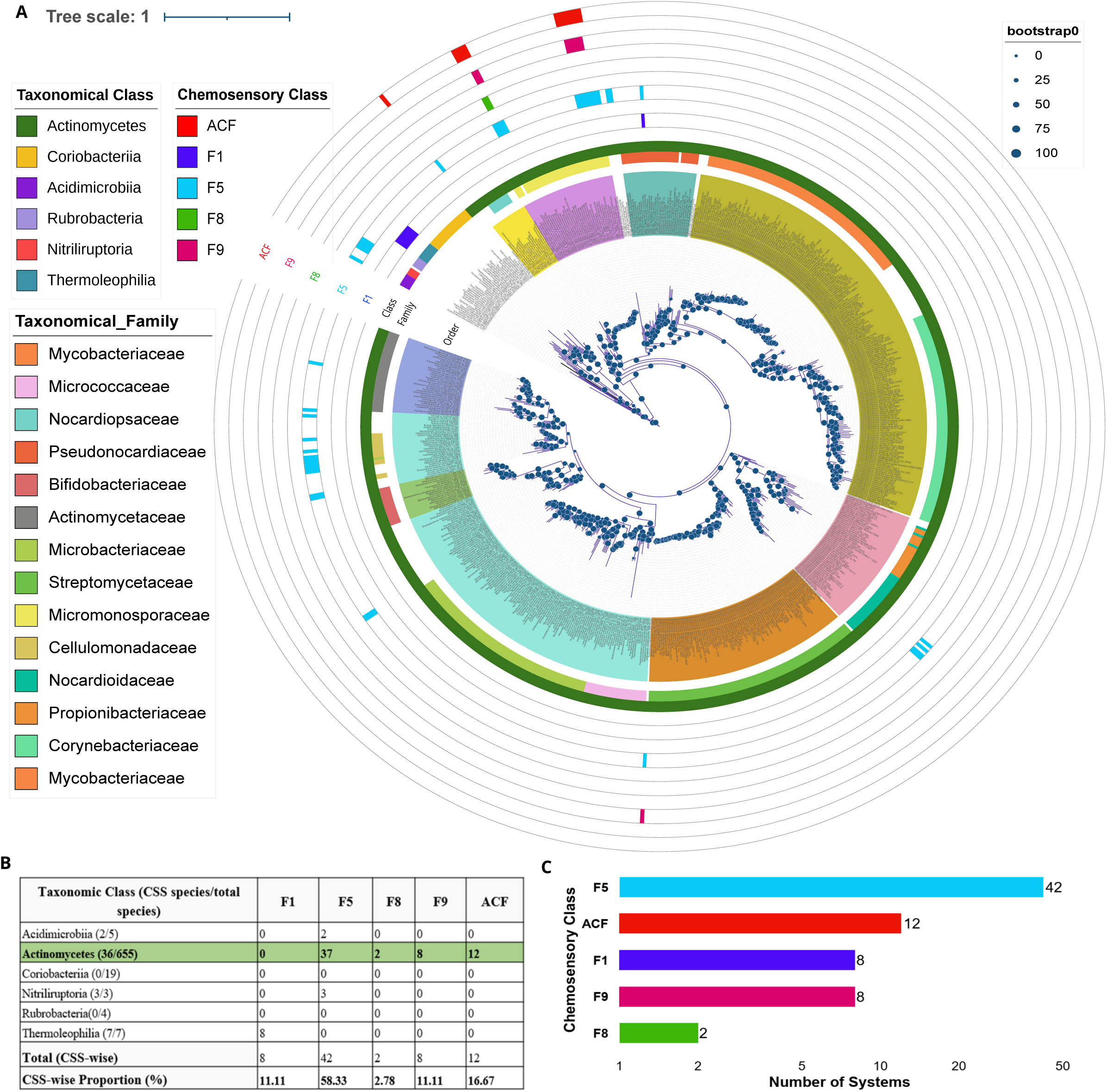
Phylogenetic distribution and abundance of chemosensory systems across Actinomycetota. **(A)** Unrooted core-protein phylogeny of 693 Actinomycetota genomes annotated with taxonomic order, class, and family followed by strips indicating the detected CSS classes. Bootstrap support values are shown as node-sized circles. The outer annotation highlights the presence and distribution of CSS classes (F1, F5, F8, F9, and ACF) across the phylogeny, with Actinomycetes forming a large, derived clade enriched in multiple CSS types. (**B)** Class-wise distribution summary of CSS counts showing the number of genomes encoding each CSS class within the Actinomycetota taxonomic classes. **(C)** Overall abundance of CSS classes across the phylum, illustrating the predominance of F5 systems, followed by ACF, F1, F9, and F8.

Further granularity at the order and genus level reveals highly constrained phylogenetic distribution of accessory CSS types. The F9 chemosensory systems were exclusively confined to the class Actinomycetes and were predominantly enriched within the family Micromonosporaceae (order Micromonosporales), where all F9-containing genomes also encoded F5 and ACF systems, indicating their strong co-occurrence and potential functional association. Notably, all examined members of the Actinoplanes (7 genomes) and Dactylosporangium (4 genomes) consistently harbored ACF architecture, whereas closely related genera including Micromonospora, Natronosporangium, Phytohabitans, Polymorphospora, and Salinispora lacked these systems entirely. This sharply partitioned distribution pattern suggests lineage-specific acquisition followed by stable evolutionary retention of these accessory modules, potentially reflecting adaptation to specialized ecological or developmental processes such as substrate colonization or sporulation. The invariant co-occurrence of F9 with both ACF and F5 systems further supports the possibility of coordinated signaling integration between multiple chemosensory pathways, consistent with the modular organization and evolutionary plasticity of bacterial signaling networks (Wuichet & Zhulin, 2010). Given that F9 systems are rarely observed in isolation (Mo et al., 2022), the persistent association of F9 with F5 in our dataset may indicate conserved inter-pathway coupling or complementary sensory functionality within these lineages.

Collectively, these findings (Fig. 3C) demonstrate that CSS evolution in Actinomycetota is highly non-uniform, characterized by early lineage simplification of F5 CSS, followed by selective expansion in sporadic organisms in Actinomycetes. The emergence of additional CSS types (F8, F9, ACF) in a few distinct lineages underscores a shift toward increased sensory resolution and ecological adaptability, reflecting the complex lifestyles and environmental interactions of these bacteria.

### Class-specific modular reorganization of chemosensory clusters accompanies the F1 to F5 transition and Actinomycetes diversification

CheA-based phylogeny of seventy-one homologs from 48 Actinomycetota genomes resolved the five CSS classes (F1, F5, F8, F9 and ACF) and showed a class-level transition across the phylum, from F1 in Thermoleophilia to F5 in the remaining lineages (Fig. 4A). This class shift is strongly reflected in changes in chemosensory system (CSS) cluster organization. Actinomycetes exhibited the greatest breadth of CSS classes and architectures, consistent with extensive system expansion and specialization. In contrast, members of the classes Nitriliruptoria and Acidimicrobia retained fewer, more conserved architectures. All seven members of Thermoleophilia displayed a distinct pattern: despite limited class diversity, they consistently harbored one to two copies of the F1 system, indicating retention and modest expansion of ancestral architecture rather than diversification into newly evolved classes. Notably, in certain genomes such as *Conexibacter woesei* and *Capillimicrobium parvum*, CheA proteins could be confidently assigned to a canonical F-class (predominantly F1), whereas associated components exhibited divergent or weak classification signals. The presence of CheA-classified systems lacking coherent supporting components suggests the existence of incomplete or hybrid chemosensory architectures, likely arising from domain shuffling and modular evolution of signaling pathways (Wuichet & Zhulin, 2010; Capra & Laub, 2012).

**Figure 4:**
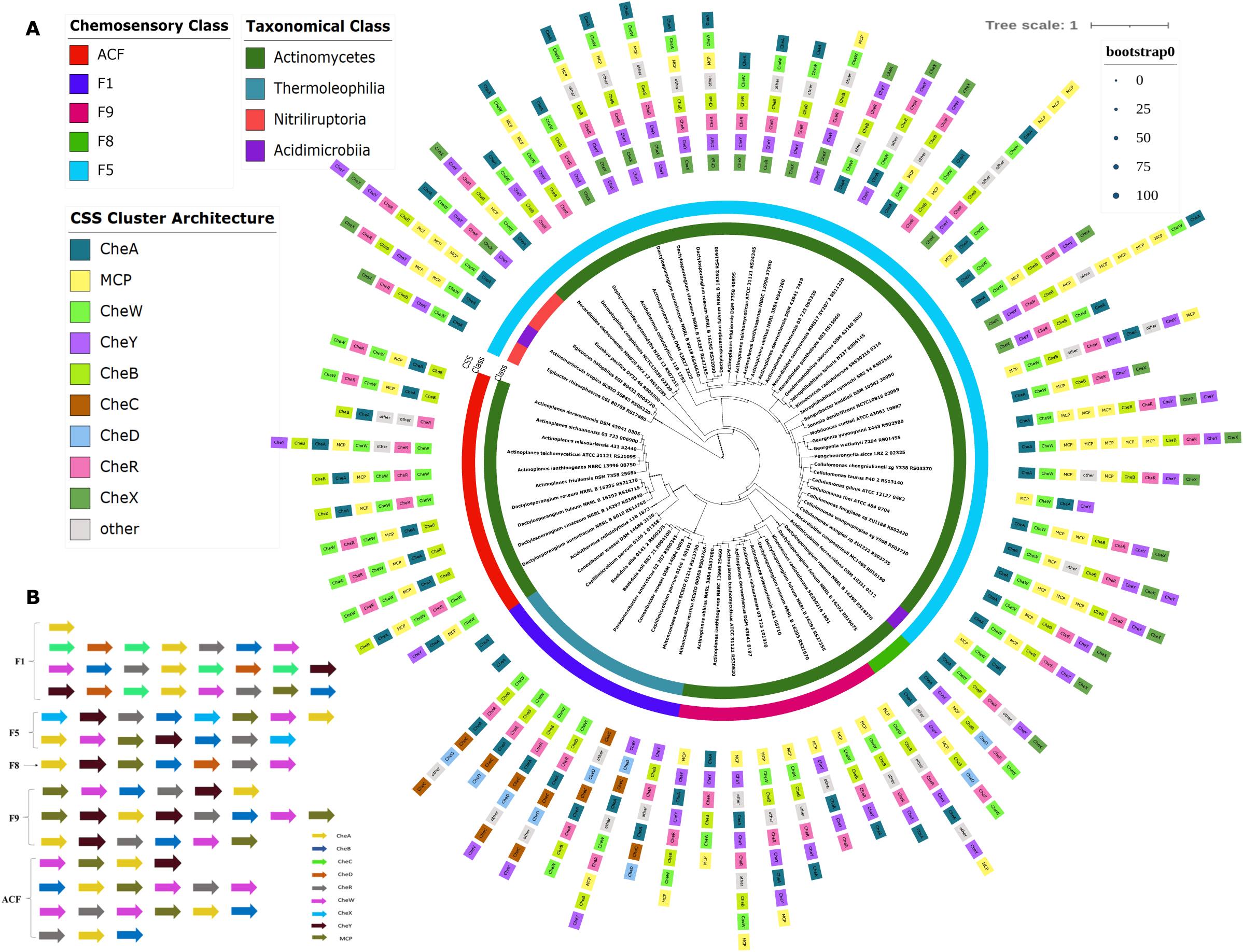
Phylogeny and modular architecture of chemosensory systems in Actinomycetota. **(A)** Unrooted circular phylogeny constructed using CheA homologs from Actinomycetota species. The inner ring indicates the corresponding taxonomical classes, and the outer ring denotes the assigned chemosensory (CSS) classes. Surrounding each tip, colored blocks represent the associated CSS cluster architecture, with individual components (CheA, CheW, CheY, CheB, CheC, CheD, CheR, CheX, MCP, and others) displayed as separate modules. Bootstrap support values are shown as proportional dot sizes near the branches. (**B)** Schematic representation of all identified CSS cluster architectures, with each arrow depicting a single protein component color-coded according to its functional module. This panel summarizes the arrangement and composition of modules across different CSS classes (F1, F5, F8, F9, and ACF).

Mapping CheA proteins to their corresponding CSS architectures further revealed distinct lineage-level modular signatures. Classes such as F5 and F8 exhibited highly reorganized CheW, CheB, and CheD modules, whereas F1 and F9 retained more canonical structural arrangements. The architectural patterns depicted in Fig. 4B highlight that each CSS class has co-evolved with characteristic combinations of accessory domains, reinforcing their functional divergence.

### Polyphyletic CheA lineages indicate multiple independent origins of Actinomycetota chemosensory systems

To resolve the evolutionary origins of Actinomycetota CSSs, we reconstructed a maximum-likelihood phylogeny of the 71 Actinomycetota CheA proteins from forty-eight genomes together with nine hundred non-redundant CheA homologs from other bacterial phyla (Fig. 5). The tree resolves five deeply branching clades that correspond to the five CSS classes (F1, F5, F8, F9 and ACF), so CheA phylogeny is congruent with class assignment based on cluster architecture. Actinomycetota sequences do not form a single clade. They fall into multiple separate lineages, each nested among homologs from other phyla, showcasing a polyphyletic distribution.

**Figure 5:**
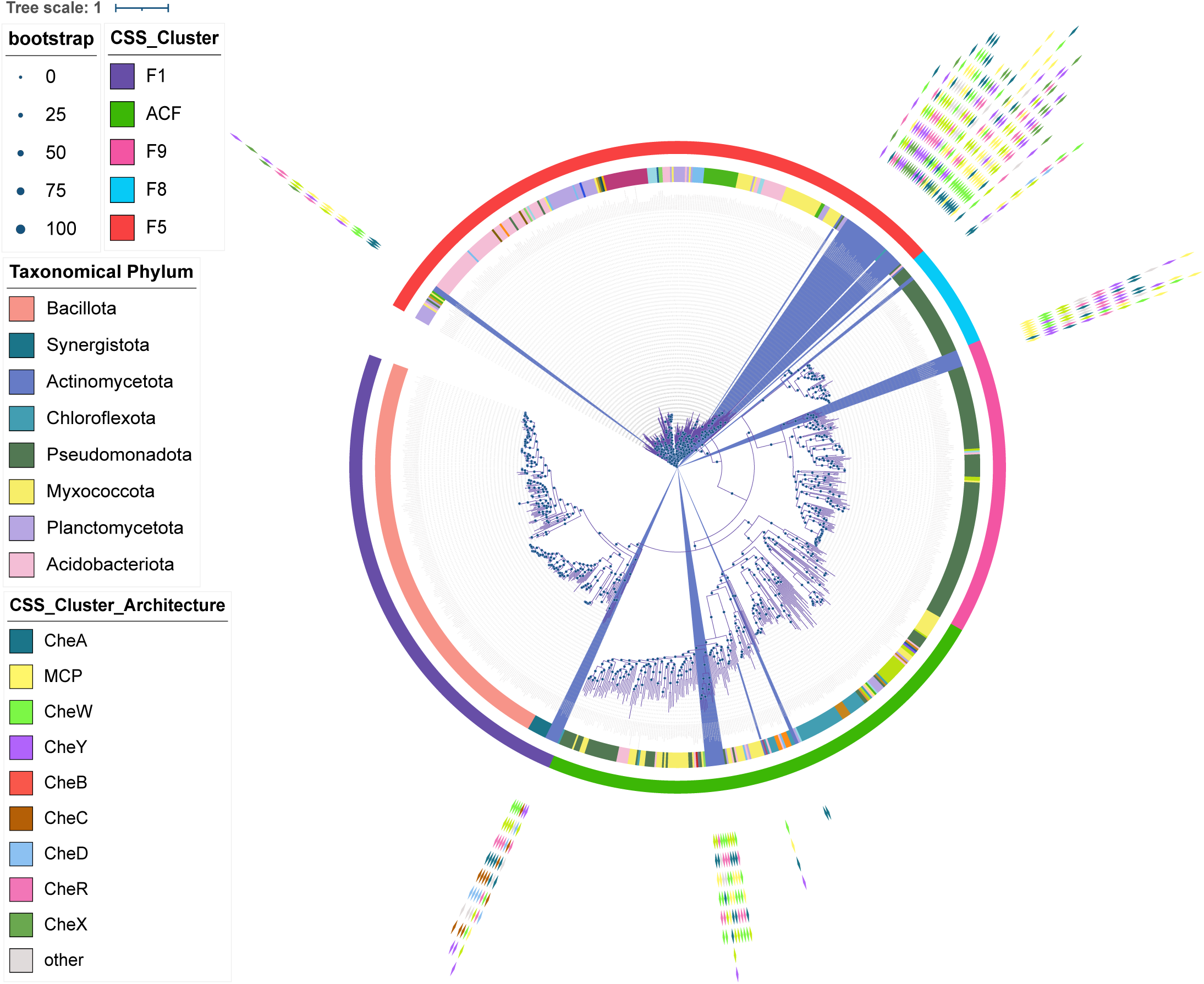
Evolutionary context of Actinomycetota CSS across bacterial phyla. Maximum-likelihood phylogeny of phylum Actinomycetota CheA and their non-redundant homologs from diverse bacterial phyla, providing a global evolutionary trajectory for CSS diversification. Actinomycetota-derived CheA sequences (highlighted clades) cluster distinctly within specific CSS lineages. Outer rings annotate CSS class (F1, F5, F8, F9, ACF) and taxonomic phylum, while inner annotations depict associated cluster architectures, including core and accessory components. This tree provides a consolidated view of the evolutionary placement, divergence, and potential horizontal acquisition of chemosensory systems within Actinomycetota relative to other bacterial lineages.

The pattern differs by class. F5 and ACF clades span many phyla, including Pseudomonadota, Myxococcota, Chloroflexota, Planctomycetota and Acidobacteriota, with Actinomycetota sequences interspersed among them. F8 and F9 form a sister grouping that is dominated by Pseudomonadota, and Actinomycetota sequences occupy discrete positions within it. The F1 clade is mainly Bacillota and Synergistota, and the Actinomycetota F1 sequences, which come from Thermoleophilia, branch within it.

Consistent with earlier observations, the presence of multiple, well-separated violet clades indicates that Actinomycetota CheA homologs do not form a single cohesive lineage. Instead, they show a polyphyletic pattern, demonstrating that these proteins have undergone lineage-specific diversification within the phylum. Such diversification is consistent with observed changes in the architecture of CSS classes. It suggests that the various CheA forms in phylum Actinomycetota did not arise from a single ancestor but rather developed independently through numerous evolutionary events. Overall, this evolutionary history reveals a complex and varied (mosaic) path for the CSS systems within the Actinomycetota phylum.

## Discussion

This high-throughput comparative genomic study of 693 Actinomycetota genomes provides newer insights into how taxonomic dominance, genome architecture, and signal-transduction evolution have shaped the diversification of this phylum. The higher availability of genomes within the Actinomycetes class likely reflects a study bias toward clinically and industrially important genera from class Actinomycetes. The representation of the class Actinomycetes, together with its broad ecological distribution and remarkable genomic flexibility (having genome size in range of 1.3 Mbp to 13 Mbp, average of 5 Mbp and median of 4.2 Mbp), highlights its central role in its evolutionary success amidst the kingdom Bacteria. The strong linear relationship between genome size and total CDS count across most lineages suggests that genome expansion in Actinomycetota is driven predominantly by increased coding capacity rather than accumulation of non-coding DNA, supporting the metabolic and ecological versatility characteristic of this phylum.

The signal transduction profile further reveals lineage-specific differences, where TCS proteins constitute a substantial fraction (25.2%), suggesting the importance of phosphorylation-based regulation in these bacteria. Chemosensory systems (CSS), however, exhibit far more restricted distribution, as only forty-eight species of four taxonomical classes encode complete CSS clusters, confirming that chemosensing is not a ubiquitous trait in Actinomycetota. Yet within non-chemosensory classes, hybrid histidine kinases are disproportionately abundant, particularly in orders *Eggerthellales* and *Rubrobacterales*, suggesting that these lineages rely on hybrid systems instead of canonical chemotaxis pathways. These patterns indicate that while the core HK–RR machinery is widespread, the evolution of dedicated chemosensory systems is both selective and lineage-dependent.

The class Actinomycetes stands out as the hotspot of CSS innovation. Basal classes such as Coriobacteriia and Rubrobacteria completely lack CSS, while members of Thermoleophilia retain only the ancestral F1 class. Interestingly, specific Thermoleophilia genomes, such as *Capillimicrobium parvum* and *Conexibacter woesei*, demonstrate atypical chemosensory system architecture. While their core signaling kinases (CheA) are phylogenetically rooted within the ancestral F1 clade, the flanking accessory proteins exhibit evolutionary divergence in operon prediction analysis, showing orthology to disparate CSS classes including ACF, F3, or F6. Such architectures likely reflect ongoing modular evolution involving operon rearrangement, selective component retention, and domain shuffling, processes previously implicated in bacterial signaling diversification (Wuichet & Zhulin, 2010; Capra & Laub, 2012). Functionally, these fragmented or hybridized architectures may preserve limited environmental sensing capacity while lacking the broader signaling plasticity associated with fully diversified CSS repertoires. Collectively, these observations position Thermoleophilia as a potentially transitional lineage representing the verge of conventional CSS reduction within Actinomycetota.

Conversely, Acidimicrobiia encodes only F5 systems, implying an early independent specialization. Actinomycetes members, however, show the higher diversity and expansion of CSS architectures. They harbor F5, F8, F9, and ACF classes indicating that this lineage is the primary driver of CSS diversification within the phylum. The dominance of F5 across the whole phylum, but not across all organisms, can thus be attributed almost entirely to its expansion within Actinomycetes. Eleven out of twelve occurrences of ACFs are present only in Micromonosporales order, which again showcases that such ACF system might have been procured from somewhere within these organisms. The modular organization of CSS further supports this diversification. The transition from ancestral F1 systems to newer architectures such as F5, F8, and F9 is marked by consistent rearrangements in accessory modules, including CheW, CheB, and CheD. These reorganizations suggest that class-specific functional innovation may have occurred through domain shuffling, module duplication, and acquisition of new interaction partners. Despite having diverse ecological habitats among order Mycobacteriales (Fig S1), their members exhibited a complete absence of chemosensory (CSS) systems across the genomes analyzed. Notably, *Mycobacterium tuberculosis,* a representative human pathogen from this order, lacks CSS pathways and flagellar machinery, consistent with its non-motile, host-adapted lifestyle as revealed by genome-scale analyses (Cole et al., 1998). Instead, environmental sensing and adaptive responses are primarily mediated through specialized two-component regulatory systems, including DosRST and other sensor kinase–response regulator pairs, which enable the detection of hypoxia, redox stress, and host-derived signals (Shiloh et al., 2008). This regulatory strategy reflects a shift from motility-driven environmental navigation to intracellular survival mechanisms governed by signal transduction and transcriptional regulation. Given the complete absence of canonical CSS pathways, it remains an open question whether additional, non-canonical sensory mechanisms such as one-component regulators, redox-sensitive transcription factors, or ligand-binding regulatory proteins contribute to environmental perception and signal integration in these organisms. It must be noted that in terms of CSS diversity and functions, another soil-residing phylum Myxococcota (Myxobacteria), which also has large genomes and complex developmental cycles like Actinomycetes, showcase the CSS diversity quite contrary to Actinomycetes. Those organisms encode up to fourteen types of CSS, with eight in *Myxococcus xanthus* DK1622, most of them tied to motility and development (Barry et al., 2006; Sharma et al., 2018).

In our study, F9 cluster is majorly present in the organisms having F5 and ACF CSS clusters within the Actinomycetes class. This co-occurrence suggests that F9 may function in coordination with other chemosensory pathways rather than operating independently. Such an arrangement could reflect functional specialization, where F9 complements other CSS clusters by sensing distinct environmental cues (Zhu et al., 2023). In *Actinoplanes* and *Dactylosporangium*, the F9 CSS cluster is likely dedicated to the specialized navigation of zoospores. While F5 or F8 clusters may manage general physiological responses or slower vegetative transitions, the F9 system appears specifically tailored to handle the high-speed chemotaxis (135–160 µm/s) characteristic of these genera (Zhu et al., 2023).

The presence of F9 alongside ACF systems suggests a sophisticated ‘search-and-arrest’ mechanism (Zhu et al., 2023). In this model, F9 detects specific nutrient gradients in aquatic environments to guide the zoospore, while the associated clusters facilitate the rapid flagellar rotation arrest required for the zoospore to settle and initiate mycelial growth in *Actinoplanes missouriensis* (Kato et al., 2024). This specialized signaling repertoire enables these organisms to thrive in dynamic environments by maximizing the efficiency of their brief motile stage (Zhu et al., 2023).

Finally, the phylum-wide CheA phylogeny created using the close homologs from NR database underscores the polyphyletic nature of Actinomycetota chemosensory evolution. Actinomycetota CheA proteins do not form a single monophyletic clade; instead, they are dispersed across several deep branches alongside homologs from diverse bacterial phyla. This polyphyly, when combined with the class-specific architectural patterns, suggests that CSS expansion in Actinomycetota did not result from a single ancestral system but rather from multiple independent evolutionary events. Horizontal gene transfer, differential gene loss, and lineage-specific innovation are likely contributors to this mosaic evolutionary pattern.

Overall, this study reveals that the Actinomycetota phylum exhibits striking heterogeneity in genomic and signaling features. While genome size expansion is universally tied to coding capacity, sensory system evolution, especially CSS diversification, is highly lineage-specific, with Actinomycetes emerging as the primary hub of innovation. These findings enhance our understanding of how ecological pressures, modular evolution, and phylogenetic history collectively drive functional diversification in bacterial phyla.

## Conclusion

This phylum-wide study using 693 complete genomes reveals that chemosensory system (CSS) evolution in phylum Actinomycetota is highly heterogeneous, with canonical systems restricted to a few selected lineages rather than conserved across the phylum. The predominance of two-component systems, alongside the enrichment of hybrid kinases in CSS-deficient lineages, highlights the diversity of regulatory architectures associated with environmental sensing. CSS diversity is concentrated in the class Actinomycetes, where F5, F8, F9, and ACF systems exhibit lineage-specific distributions and distinct cluster architectures. The co-occurrence of multiple CSS classes and the reorganization of accessory components suggest that modular changes have contributed to their diversification. Furthermore, the polyphyletic distribution of Actinomycetota CheA homologs across bacterial phyla supports a mosaic evolutionary history potentially involving horizontal gene transfer, differential gene loss, and independent lineage-specific diversification. These findings argue against a single, uniform trajectory of CSS evolution and instead support a model of selective acquisition, retention, and architectural remodeling across Actinomycetota. Future phylogenetic reconciliation and functional characterization of divergent CSS architectures will be essential to distinguish these evolutionary processes and establish how individual systems contribute to the ecological and physiological adaptations of their hosts.

## Supplementary data legends

**Figure S1. Heatmap illustrates the ecological distribution of 693 organisms across taxonomic orders**, with cell values representing the number of species associated with each of the four designated ecological niches: host-associated, sea, terrestrial land, and terrestrial water. Ecological annotations were compiled from published literature and curated databases, including BacDive, for approximately 350 genomes with available habitat information.

**Figure S2. Box plot illustrates the distribution of genome size (Mbp) across taxonomic orders within five bacterial classes.** The width of each box reflects the kernel density estimate of the data, capturing the full range and frequency distribution of genome sizes within each order.

**Figure S3. Box plot represents the variation in GC content (%) across taxonomic orders within the five major bacterial classes examined.** The distribution profiles highlight inter- and intra-order variability in nucleotide composition.

**Figure S4. Scatter-density plot examining the relationship between genome size (Mbp) and GC content (%) across taxonomic orders (indicated in the legend).** The overall distribution reveals a weak positive correlation (Pearson’s r = 0.5127), suggesting a modest but statistically notable association between genomic size and nucleotide composition.

**Figure S5. Scatter-density plot depicting the relationship between genome size (Mbp) and total coding sequence (CDS) count across taxonomic orders.** A strong positive correlation (Pearson’s r = 0.9367) is observed, indicating that CDS count scales robustly with genome size across the taxonomic diversity examined.

## Supplementary Tables

**Table S1:** Taxonomic and Genomic Metadata for Actinomycetota Genomes. Table showing the distribution of genome size, GC content, protein coding gene count, ecology, taxonomy, and NCBI assembly accession among Actinomycetota isolates.

**Table S2:** Distribution of TCS constituent proteins and CheA across Actinomycetota genomes: Table depicts the quantitative distribution of TCS components like orphan histidine kinases, orphan response regulators, hybrid components (HHK, HRR) and the primary CSS component, CheA among Actinomycetota isolates.

**Table S3:** Distribution of CSS functional categories across Actinomycetota genomes: Table shows the quantitative distribution of chemosensory system (CSS) categories, including F1, F5, F8, F9, and ACF pathways, across Actinomycetota genomes. Values represent the copy number of each CSS class detected per genome along with their taxonomical classification.

## Data availability statement

Publicly available datasets were analyzed in this study. The NCBI accession numbers of these datasets are available in “G” column in supplementary table S1.

## Author contributions

SNS: Investigation, Methodology, Resources, Validation, Writing – original draft, Data curation, Formal analysis, Visualization.

SB: Validation, Writing – original draft, Data curation, Formal analysis, Visualization.

GS: Methodology, Resources, Validation, Writing – review & editing, Conceptualization, Funding acquisition, Project administration, Supervision.

## Acknowledgments

The support and computational resources provided by PARAM Seva Facility under the National Supercomputing Mission (NSM), Government of India at the Indian Institute of Technology Hyderabad are gratefully acknowledged.

## Reference

1. Krell, T., Lacal, J., Busch, A., Silva-Jiménez, H., Guazzaroni, M. E., & Ramos, J. L. (2010). Bacterial sensor kinases: diversity in the recognition of environmental signals. Annual review of microbiology, 64, 539–559. 10.1146/annurev.micro.112408.134054

2. Ulrich, L. E., Koonin, E. V., & Zhulin, I. B. (2005). One-component systems dominate signal transduction in prokaryotes. Trends in microbiology, 13(2), 52–56. 10.1016/j.tim.2004.12.006.

3. Galperin, M. Y., Higdon, R., & Kolker, E. (2010). Interplay of heritage and habitat in the distribution of bacterial signal transduction systems. Molecular BioSystems, 6(4), 721–728. 10.1039/B908047C.

4. Stock, A. M., Robinson, V. L., & Goudreau, P. N. (2000). Two-component signal transduction. Annual review of biochemistry, 69, 183–215. 10.1146/annurev.biochem.69.1.183

5. Gao, R., & Stock, A. M. (2009). Biological insights from structures of two-component proteins. Annual review of microbiology, 63, 133–154. 10.1146/annurev.micro.091208.073214

6. Bi, S., & Sourjik, V. (2018). Stimulus sensing and signal processing in bacterial chemotaxis. Current opinion in microbiology, 45, 22–29. 10.1016/j.mib.2018.02.002

7. Sourjik, V., & Berg, H. C. (2002). Binding of the Escherichia coli response regulator CheY to its target measured in vivo by fluorescence resonance energy transfer. Proceedings of the National Academy of Sciences of the United States of America, 99(20), 12669–12674. 10.1073/pnas.192463199

8. Hazelbauer, G. L., Falke, J. J., & Parkinson, J. S. (2008). Bacterial chemoreceptors: high-performance signaling in networked arrays. Trends in biochemical sciences, 33(1), 9–19. 10.1016/j.tibs.2007.09.014

9. Stock, J. B., & Koshland, D. E. (1978). A protein methylesterase involved in bacterial chemotaxis. Proceedings of the National Academy of Sciences, 75(8), 3659–3663. 10.1073/pnas.75.8.3659

10. Kentner, D., & Sourjik, V. (2006). Spatial organization of the bacterial chemotaxis system. Current opinion in microbiology, 9(6), 619–624. 10.1016/j.mib.2006.10.012

11. Briegel, A., Li, X., Bilwes, A. M., Hughes, K. T., Jensen, G. J., & Crane, B. R. (2012). Bacterial chemoreceptor arrays are hexagonally packed trimers of receptor dimers interconnected by rings of CheA and CheW. Proceedings of the National Academy of Sciences, 109(10), 3766–3771. 10.1073/pnas.1115719109

12. Cassidy, C. K., Himes, B. A., Sun, D., Ma, J., Zhao, G., Parkinson, J. S., Schulten, K., & Zhang, P. (2020). Structure and dynamics of the *E. coli* chemotaxis core signaling complex by cryo-electron tomography and molecular simulations. Commun Biol 3, 24. 10.1038/s42003-019-0748-0

13. Zhao, R., Collins, E. J., Bourret, R. B., & Silversmith, R. E. (2002). Structure and mechanism of Action of a two-component system phosphatase: CheZ. Nature Structural Biology, 9(8), 570–575. 10.1038/nsb816

14. He, K., & Bauer, C. E. (2014). Chemosensory signaling systems that control bacterial survival. In Trends in Microbiology (Vol. 22, Issue 7, pp. 389–398). Elsevier Ltd. 10.1016/j.tim.2014.04.004

15. Munar-Palmer, M., Santamaría-Hernando, S., Liedtke, J., Ortega, D. R., López-Torrejón, G., Rodríguez-Herva, J. J., Briegel, A., & López-Solanilla, E. (2024). Chemosensory systems interact to shape relevant traits for bacterial plant pathogenesis. mBio, 15(7), e0087124. 10.1128/mbio.00871-24

16. Colin, R., Ni, B., Laganenka, L., & Sourjik, V. (2021). Multiple functions of flagellar motility and chemotaxis in bacterial physiology. FEMS microbiology reviews, 45(6), fuab038. 10.1093/femsre/fuab038

17. Wuichet, K., & Zhulin, I. B. (2010). Origins and diversification of a complex signal transduction system in prokaryotes. Science Signaling, 3(128). 10.1126/scisignal.2000724

18. Zhulin I. B. (2001). The superfamily of chemotaxis transducers: from physiology to genomics and back. Advances in microbial physiology, 45, 157–198. 10.1016/s0065-2911(01)45004-1

19. Sharma, G., Khatri, I., & Subramanian, S. (2018). Comparative Genomics of Myxobacterial Chemosensory Systems, 200:10.1128/jb.00620-17. https://doi.org/10.1128/jb.00620-17

20. Alexander, R. P., & Zhulin, I. B. (2007). Evolutionary genomics reveals conserved structural determinants of signaling and adaptation in microbial chemoreceptors. Proceedings of the National Academy of Sciences of the United States of America, 104(8), 2885–2890. 10.1073/pnas.0609359104

21. Seshadri, R., Roux, S., Huber, K. J., Wu, D., Yu, S., Udwary, D., Call, L., Nayfach, S., Hahnke, R.L., Pukall, R.,..& Ivanova, N. N. (2022). Expanding the genomic encyclopedia of Actinobacteria with 824 isolate reference genomes. Cell Genomics, 2(12). 10.1016/j.xgen.2022.100213

22. Hashim, H. S., Zayan, M. M., Mohamed, A. A., & El Rahman Abulila, H. I. A. (2026). Actinomycetes in the spotlight: biodiversity and their role in bioremediation. World journal of microbiology & biotechnology, 42(2), 44. 10.1007/s11274-025-04610-5

23. Barka, E. A., Vatsa, P., Sanchez, L., Gaveau-Vaillant, N., Jacquard, C., Meier-Kolthoff, J. P., Klenk, H. P., Clément, C., Ouhdouch, Y., & van Wezel, G. P. (2016). Taxonomy, Physiology, and Natural Products of Actinobacteria. Microbiology and molecular biology reviews : MMBR, 80(1), 1–43. 10.1128/MMBR.00019-15

24. Bentley, S. D., Chater, K. F., Cerdeño-Tárraga, A. M., Challis, G. L., Thomson, N. R., James, K. D., Harris, D. E., Quail, M. A., Kieser, H., Harper, D., Bateman, A., Brown, S., Chandra, G., Chen, C. W., Collins, M., Cronin, A., Fraser, A., Goble, A., Hidalgo, J., Hornsby, T., Howarth, S., Huang, C.-H., Kieser, T., Larke, L., Murphy, L., Oliver, K., O’Neil, S., Rabbinowitsch, E., Rajandream, M.-A., Rutherford, K., Rutter, S., Seeger, K., Saunders, D., Sharp, S., Squares, R., Squares, S., Taylor, K., Warren, T., Wietzorrek, A., Woodward, J., Barrell, B. G., Parkhill, J., & Hopwood, D. A. (2002). Complete genome sequence of the model actinomycete *Streptomyces coelicolor* A3(2). Nature, 417(6885), 141–147. 10.1038/417141a

25. Jorgensen, T. S., Mohite, O. S., Sterndorff, E. B., Alvarez-Arevalo, M., Blin, K., Booth, T. J., Charusanti, P., Faurdal, D., Hansen, T., Nuhamunada, M., Mourched, A. S., Palsson, B., & Weber, T. (2024). A treasure trove of 1034 actinomycete genomes. Nucleic Acids Research, 52(13), 7487–7503. 10.1093/nar/gkae523

26. Antoraz, S., Santamaría, R. I., Díaz, M., Sanz, D., & Rodríguez, H. (2020). Toward a new focus in antibiotic and drug discovery from the Streptomyces arsenal. Scientific Reports, 10, 8573. 10.1038/s41598-020-5890

27. Cuervo, L., Méndez, C., Salas, J. A., Olano, C., & Malmierca, M. G. (2024). Volatile communication in Actinobacteria: a language for secondary metabolism regulation. Microbial Cell Factories, 23, 177. 10.1186/s12934-024-02456-4

28. Wadach, P., Jakimowicz, D., & Gongerowska-Jac, M. (2025). Untangling the Complexity of Two-Component Signal Transduction in Bacteria. Microorganisms, 13(9), 2013. 10.3390/microorganisms13092013

29. Sanchez de la Nieta, R., Santamaría, R. I., & Díaz, M. (2022). Two-Component Systems of *Streptomyces coelicolor*: An Intricate Network to Be Unraveled. International journal of molecular sciences, 23(23), 15085. 10.3390/ijms232315085

30. Hutchings, M. I., Hoskisson, P. A., Chandra, G., & Buttner, M. J. (2004). Sensing and responding to diverse extracellular signals? Analysis of the sensor kinases and response regulators of *Streptomyces coelicolor* A3(2). Microbiology, 150(9), 2795–2806. 10.1099/mic.0.27181-0

31. Paget, M. S., Hong, H. J., Bibb, M. A., & Buttner, M. J. (2002). The ECF sigma factors of *Streptomyces coelicolor* A3(2). Journal of Molecular Microbiology and Biotechnology, 4(4), 417–422.

32. McLean TC, Beaton ADM, Holmes NA, Martins C, Saalbach G, Chandra G, Batey SFD, China J, Wilkinson B, Hutchings MI.2026.The conserved two-component systems CutRS and CssRS control the protein secretion stress response in Streptomyces. mBio17:e02991–25.10.1128/mbio.02991-25

33. Cruz-Bautista, R., Ruíz-Villafán, B., Romero-Rodríguez, A., Rodríguez-Sanoja, R., & Sánchez, S. (2023). Trends in the two-component system’s role in the synthesis of antibiotics by Streptomyces. Applied microbiology and biotechnology, 107(15), 4727–4743. 10.1007/s00253-023-12623-z

34. Gongerowska-Jac M, Szafran MJ, Mikołajczyk J, Szymczak J, Bartyńska M, Gierlikowska A, Biały S, Elliot MA, Jakimowicz D. (2021).Global Chromosome Topology and the Two-Component Systems in Concerted Manner Regulate TranscriptioninStreptomyces. mSystems6:e01142–21.10.1128/mSystems.01142-2.

35. Muñoz-Dorado, J., Marcos-Torres, F. J., García-Bravo, E., Moraleda-Muñoz, A., & Pérez, J. (2016). Myxobacteria: Moving, killing, feeding, and surviving together. In Frontiers in Microbiology (Vol. 7, Issue MAY). Frontiers Media S.A. 10.3389/fmicb.2016.00781

36. Shi, X., Wegener-Feldbrügge, S., Huntley, S., Hamann, N., Hedderich, R., & Søgaard-Andersen, L. (2008). Bioinformatics and experimental analysis of proteins of two-component systems in Myxococcus xanthus. Journal of bacteriology, 190(2), 613–624. 10.1128/JB.01502-07

37. Barry S. Goldman, W. C. Nierman, Dale Kaiser, et al. (2006). Evolution of sensory complexity recorded in a myxobacterial genome. Proceedings of the National Academy of Sciences of the United States of America, 103(41), 15200–15205. 10.1073/pnas.0607335103

38. Anik, T. A. (2025). FetchM: Streamlining Genome and Metadata Integration for Microbial Comparative Genomics. 10.1101/2025.04.08.647722

39. Mistry, J., Chuguransky, S., Williams, L., Qureshi, M., Salazar, G. A., Sonnhammer, E. L. L., Tosatto, S. C. E., Paladin, L., Raj, S., Richardson, L. J., Finn, R. D., & Bateman, A. (2021). Pfam: The protein families database in 2021. Nucleic acids research, 49(D1), D412–D419. 10.1093/nar/gkaa913

40. Finn, R. D., Clements, J., & Eddy, S. R. (2011). HMMER web server: interactive sequence similarity searching. Nucleic acids research, 39(Web Server issue), W29–W37. 10.1093/nar/gkr367

41. Gumerov, V. M., Ulrich, L. E., & Zhulin, I. B. (2024). MiST 4.0: a new release of the microbial signal transduction database, now with a metagenomic component. Nucleic Acids Research, 52(D1), D647–D653. 10.1093/nar/gkad847

42. Lechner, M., Findeiss, S., Steiner, L., Marz, M., Stadler, P. F., & Prohaska, S. J. (2011). Proteinortho: detection of (co-)orthologs in large-scale analysis. BMC bioinformatics, 12, 124. 10.1186/1471-2105-12-124

43. Edgar R. C. (2022). Muscle5: High-accuracy alignment ensembles enable unbiased assessments of sequence homology and phylogeny. Nature communications, 13(1), 6968. 10.1038/s41467-022-34630-w

44. Price, M. N., Dehal, P. S., & Arkin, A. P. (2010). FastTree 2—approximately maximum-likelihood trees for large alignments. PLOS ONE, 5(3), e9490. 10.1371/journal.pone.0009490

45. Letunic, I., & Bork, P. (2024). Interactive Tree of Life (iTOL) v6: recent updates to the phylogenetic tree display and annotation tool. Nucleic Acids Research, 52(W1), W78–W82. 10.1093/nar/gkae268

46. Camacho, C., Coulouris, G., Avagyan, V., Ma, N., Papadopoulos, J., Bealer, K., & Madden, T. L. (2009). BLAST+: architecture and applications. BMC bioinformatics, 10, 421. 10.1186/1471-2105-10-421

47. Sayers, E. W., Beck, J., Bolton, E. E., Brister, J. R., Chan, J., Connor, R., Feldgarden, M., Fine, A. M., Funk, K., Hoffman, J., Kannan, S., Kelly, C., Klimke, W., Kim, S., Lathrop, S., Marchler-Bauer, A., Murphy, T. D., O’Sullivan, C., Schmieder, E., Skripchenko, Y., … Pruitt, K. D. (2025). Database resources of the National Center for Biotechnology Information in 2025. Nucleic acids research, 53(D1), D20–D29. 10.1093/nar/gkae979

48. Toft, C., & Andersson, S. G. (2010). Evolutionary microbial genomics: insights into bacterial host adaptation. Nature reviews. Genetics, 11(7), 465–475. 10.1038/nrg2798

49. Hildebrand F, Meyer A, Eyre-Walker A (2010) Evidence of Selection upon Genomic GC-Content in Bacteria. PLOS Genetics 6(9): e1001107. 10.1371/journal.pgen.1001107

50. Bohlin, J., Snipen, L., Hardy, S. P., Kristoffersen, A. B., Lagesen, K., Dønsvik, T., Skjerve, E., & Ussery, D. W. (2010). Analysis of intra-genomic GC content homogeneity within prokaryotes. BMC Genomics, 11(1), 464. 10.1186/1471-2164-11-464

51. Foerstner, K. U., von Mering, C., Hooper, S. D., & Bork, P. (2005). Environments shape the nucleotide composition of genomes. EMBO Reports, 6(12), 1208–1213. 10.1038/sj.embor.7400538

52. Cimermancic, P., Medema, M. H., Claesen, J., Kurita, K., Wieland Brown, L. C., Mavrommatis, K., Pati, A., Godfrey, P. A., Koehrsen, M., Clardy, J., Birren, B. W., Takano, E., Sali, A., Linington, R. G., & Fischbach, M. A. (2014). Insights into secondary metabolism from a global analysis of prokaryotic biosynthetic gene clusters. Cell, 158(2), 412–421. 10.1016/j.cell.2014.06.034

53. Huang, X., Pinto, D., Fritz, G., & Mascher, T. (2015). Environmental sensing in Actinobacteria: A comprehensive survey on the signaling capacity of this phylum. Journal of Bacteriology, 197(15), 2517–2535. 10.1128/JB.00176-15

54. Laub, M. T., & Goulian, M. (2007). Specificity in two-component signal transduction pathways. Annual review of genetics, 41, 121–145. 10.1146/annurev.genet.41.042007.170548

55. Groban, E. S., Clarke, E. J., Salis, H., & Voigt, C. A. (2009). Kinetic buffering of cross talk between bacterial two-component sensors. Journal of Molecular Biology, 390(3), 380–393. 10.1016/j.jmb.2009.05.007

56. Capra, E. J., & Laub, M. T. (2012). Evolution of two-component signal transduction systems. Annual review of microbiology, 66(1), 325–347. 10.1146/annurev-micro-092611-150039

57. Mo, R., Liu, Y., Chen, Y., Mao, Y., & Gao, B. (2022). Evolutionary principles of bacterial signaling capacity and complexity. mBio, 13(3), e00764–22.. 10.1128/mbio.00764-22

58. Moran N. A. (2002). Microbial minimalism: genome reduction in bacterial pathogens. Cell, 108(5), 583–586. 10.1016/s0092-8674(02)00665-7

59. Giovannoni, S. J., Cameron, J. C., & Temperton, B. (2014). Implications of streamlining theory for microbial ecology. The ISME Journal, 8(8), 1553–1565. 10.1038/ismej.2014.60

60. Shiloh, M. U., Manzanillo, P., & Cox, J. S. (2008). *Mycobacterium tuberculosis* senses host-derived carbon monoxide during macrophage infection. Cell Host & Microbe, 3(5), 323–330. 10.1016/j.chom.2008.03.007

61. Cole, S. T., Brosch, R., Parkhill, J., Garnier, T., Churcher, C., Harris, D., Gordon, S. V., Eiglmeier, K., Gas, S., Barry, C. E., 3rd, Tekaia, F., Badcock, K., Basham, D., Brown, D., Chillingworth, T., Connor, R., Davies, R., Devlin, K., Feltwell, T., Gentles, S., … Barrell, B. G. (1998). Deciphering the biology of Mycobacterium tuberculosis from the complete genome sequence. Nature, 393(6685), 537–544. 10.1038/31159

62. Zhu, S., Sun, X., Li, Y., Feng, X., & Gao, B. (2023). The common origin and degenerative evolution of flagella in Actinobacteria. Mbio, 14(6), e02526–23. 10.1128/mbio.02526-23

63. Kato, H., Tanemura, H., Kimura, T., Katsuyama, Y., Tezuka, T., & Ohnishi, Y. (2024). Molecular mechanism of flagellar motor rotation arrest in bacterial zoospores of Actinoplanes missouriensis before germination. Communications biology, 7(1), 1405. 10.1038/s42003-024-07104-6.

